# Characterizing the Impact of Nucleoid-Associated Proteins on HU-DNA Interactions by Live-Cell Single-Molecule Tracking

**DOI:** 10.64898/2025.12.19.695591

**Authors:** D. E. H. Fuller, X. Dai, L. A. McCarthy, L. E. Way, X. Wang, J. S. Biteen

## Abstract

The bacterial nucleoid undergoes extensive structural reorganization during growth, influenced by nucleoid-associated proteins (NAPs) whose interactions and effects on nucleoid organization remain unclear. We investigated these interactions by tracking single molecules of the NAP HUα-PAmCherry in living *Escherichia coli* cells in different growth phases, and we further examined how two NAPs, Dps and H-NS, impact HUα dynamics. HUα mobility varies with growth phase: In exponential phase, HUα has two distinct mobility states: a fast-diffusing state and a slower, interacting state. In stationary phase, we observed a third population of very slow molecules, suggesting stable HUα binding or confinement within compacted DNA. Deleting *dps* increases HUα mobility in stationary phase, consistent with findings that Dps promotes short-range DNA contacts and nucleoid compaction in deep stationary phase. We measured in exponential phase that *hns* deletion leads to nucleoid compaction, faster HUα diffusion, and a third population of very slow HUα molecules in these cells. In stationary phase, deleting *hns* increases these stably bound HUα molecules. Our results show that growth-phase-dependent nucleoid reorganization by Dps and H-NS influences the behavior and function of other NAPs.

**Graphical Abstract:** 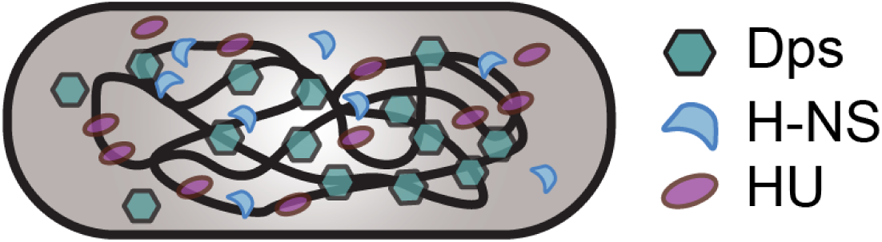

The nucleoid-associated proteins Dps, H-NS, and HU shape the bacterial chromosome in the deep stationary phase through their interactions with the nucleoid.

## Introduction

Bacteria are subject to diverse environmental conditions and frequently adapt to stress, and this response requires the protection and organization of genomic DNA. The bacterial nucleoid comprises the chromosome and DNA-binding proteins. Although it lacks a protective membrane, the nucleoid is highly organized.^1^ Mechanisms for nucleoid compartmentalization include macromolecular crowding, charge neutralization, and mechanical stress from DNA supercoiling and nucleoid proteins.^2–4^ More recently, NAPs have been recognized as drivers of both rapid response to environmental stress and of nucleoid organization, including by phase separation.^5–8^ A part of a larger stress response system,, NAPs upregulate the expression of stress response proteins,^8,9^ compact the nucleoid to physically shield genomic DNA,^10^ and catalytically neutralize perturbations.^11,12^

The DNA-binding protein from starved cells (Dps) is highly conserved among bacteria^13^ and helps to protect DNA from stressors such as oxidative damage, UV radiation, pH imbalance, and starvation.^10,11,13–15^ It has long been accepted that when nutrients are depleted by starvation, bacteria enter the stationary phase and activate stress-resistance mechanisms to maintain the nucleoid structure for cell survival.^16,17^ Additionally, recent studies have shown that energy-spilling mechanisms such as overflow metabolism and futile cycles strongly influence the metabolic pathways that are coupled to stationary-phase physiology.^18^ Dps is the most abundant NAP in stationary-phase cells, with ∼180,000 copies per cell.^19,20^ In addition to the chemical protection provided by the ferritin-like Dps protein,^11,21,22^ Dps responds to stress by non-specifically binding DNA,^23^ and, in stationary phase, Dps facilitates short-range DNA-DNA interactions.^24^ While the function of Dps has been well characterized *in vitro*, how Dps mediates nucleoid morphology *in vivo* is not fully understood.^24^ On one hand, Dps-mediated nucleoid compaction in deep stationary phase has been found not to affect RNA polymerase (RNAP) activity.^24^ Thus, the stationary-phase nucleoid remains dynamic and accessible to protein-DNA interactions. On the other hand, electron microscopy images of fixed, stationary-phase *E. coli* cells have shown highly organized crystalline arrays of Dps and DNA, indicating that Dps can organize the nucleoid into a hexagonally packed biocrystal.^23^

The histone-like nucleoid structure protein (H-NS)^25^ also plays major roles in gene regulation and nucleoid organization.^26–29^ H-NS (∼20,000 copies/cell)^19^ is a global gene regulator in *E. coli*, directly or indirectly affecting the expression of approximately 5% of genes in the *E. coli* genome,^30^ many of which are involved in adaptation to changes in environmental conditions.^31^ H-NS forms dimers that can assemble into higher-order structures to facilitate organization.^32,33^ For instance, H-NS promotes the formation of DNA loops in the nucleoid and can bridge DNA.^27^ Furthermore, in exponential phase, H-NS induces hairpin-like structures that can spatially interact, creating clusters of silenced regions in non-transcribed regions of the chromosome.^34^ H-NS has been recently reported to also impact cells in stationary phase, despite the lower concentration of H-NS (∼10,000 copies/cell).^19,35^ We recently found that H-NS mediates long-range DNA looping and is enriched at DNA loop anchors, which correlates with enhanced gene silencing in stationary phase, and H-NS reallocates transcriptional resources in stationary phase to reshape transcription.^35^

Here, we assessed how the NAPs Dps and H-NS affect the *E. coli* nucleoid structure and microenvironment based on tracking and super-resolving a fusion of the NAP HUα, which interacts non-specifically with DNA,^36^ to the photoactivatable fluorescent protein PAmCherry. For each single-molecule HUα-PAmCherry trajectory, we inferred the number of underlying mobility states, the average diffusion coefficient of single molecules in each state, the fraction of molecules in each state at a given time, and the probability of transitioning from one state to another based on the nonparametric Bayesian statistics algorithm SMAUG.^37^ The motion of HUα-PAmCherry indicates its interactions with the chromosome: immobile HUα-PAmCherry is DNA-bound, slow HUα-PAmCherry molecules are diffusing more slowly because of interactions with the nucleoid, and quickly diffusing HUα-PAmCherry molecules are free within the cell.^38^ Because the nucleoid compaction we measure is on the whole nucleoid scale, it doesn’t directly measure the reorganization of the nucleoid on the nanometer scale. We expect that the dynamics of HUα are more sensitive to nanometer-scale nucleoid regions of DNA that we cannot entirely capture within our nucleoid compaction assays. We measured the effects of Dps and H-NS on the nucleoid by tracking single HUα-PAmCherry molecules in wild-type (WT), Δ*dps*, and Δ*hns E. coli* cells, and we determined the effect of starvation stress by comparing cells in exponential and stationary phases. Overall, we found that Dps slows HUα-PAmCherry dynamics in stationary phase, while having no measurable effect in exponential phase, as expected given the very low Dps expression levels in the exponential phase.^19^ We also found that H-NS impacts the distribution of HUα-PAmCherry dynamics in exponential phase while subtly slowing HUα-PAmCherry mobility in stationary phase. Overall, these results indicate that nucleoid organization by Dps and H-NS is growth phase-dependent and that functions of NAPs are interdependent; HUα-PAmCherry dynamics are altered by the deletion of other NAPs.

## Experimental Section

### *E. coli* Growth Conditions

All strains used in this study were derived from the K-12 strain W3110 (CGSC no. 4474)^39^ and contained *hupA::PAmCherry FRT-cam-FRT* for imaging.^36^ Cells were streaked from glycerol stocks stored at −80 °C onto LB-agar plates containing 20 µg/mL chloramphenicol (CAS no. 56-75-7), then incubated at 30 °C. Isolated single colonies were then picked and grown in High-Def Azure (HDA) medium (buffered to pH 7.2; Teknova cat. no. 3H5000) supplemented with 0.2% glucose (m/v) and grown overnight at 30 °C with shaking at 250 rpm. Precultures were then diluted 1:100 into 50 mL fresh HDA medium supplemented with 0.2% glucose (m/v) in 150-mL Erlenmeyer flasks to allow ample headroom, and incubated at 30 °C with shaking at 250 rpm. For cultures in exponential phase, samples were removed at OD_600_ = 0.30 – 0.40 (Biochrom WPA CO8000 cell density meter). Optical density measurements were done in standard cuvettes with 1-cm path length. Samples were removed at 96 h for stationary-phase experiments. At least three biological replicates were studied per strain.

### *E. coli* Strain List and Construction

Strains *cWX2782* [W3110, *hupA::PAmCherry FRT-cam-FRT*] and *cWX3017* [W3110, *Δdps FRT-kan-FRT, hupA::PAmCherry FRT-cam-FRT*] were previously described in McCarthy et al.^24^

Strain *cWX2889* [W3110, *Δhns FRT, hupA::PAmCherry FRT-cam-FRT*] was generated by P1 transduction of cWX2886[[W3110, *Δhns FRT*] ^35^ with P1 lysate from strain *hupA::PAmCherry-frt-cmR-frt*.^36^ Transductants were selected for on LB-chloramphenicol 20 µg/mL plates.

### Single-Molecule Microscopy

For each replicate, 10 mL of the cell culture was extracted at the appropriate time point and then centrifuged at 6600 × *g* for 7 minutes. The supernatant was extracted and filtered twice with 0.22 µm filters to prepare spent medium. Spent medium was then used to make 2% (m/v) agarose pads on an argon plasma-etched No. 1 glass coverslip. Before imaging, 1.5 µL of cells growing at specified time points were deposited on the agarose pad with 1.5 µL of Fluoresbrite carboxylate YG beads (0.35 µm, Polysciences) suspended in spent medium (2 µL of beads diluted into 1 mL of spent medium). We previously measured ∼2.6 billion colony-forming units/mL at 96 h for WT cells.^24^ The deposited sample was then sandwiched onto the agarose pad by an argon plasma-etched No. 1 glass coverslip.

Experiments were conducted on an Olympus IX-71 microscope with either a 100× 1.40 NA or a 100× 1.45 NA phase-contrast oil-immersion objective heated to 30 °C with an objective heater (Bioptics). Objective immersion oil optimized for 30 °C (Zeiss) was used to reduce optical aberrations. Photoactivation of HUα-PAmCherry was performed with a 405-nm laser (Coherent Cube 405-100) with a 1 W/cm^2^ power density. The pulse duration of the 405-nm laser was 100 – 300 ms. Imaging of HUα-PAmCherry was carried out with a 561-nm laser (Coherent Sapphire 561-50) with a power density of 0.46 kW/cm^2^. The fluorescence emission of HUα-PAmCherry was filtered with a 561-nm long-pass filter. Images were collected with a 512 × 512-pixel Photometrics Evolve electron-multiplying charge-coupled device camera. A 40-ms exposure time was used for tracking. Sample drift was monitored with fluorescent beads located near the cells. Each cell was imaged for no longer than 1 h.

### Single-Molecule Microscopy Analysis

#### Single-Molecule Trajectory Analysis

Cell segmentation from phase-contrast images was performed with the Omnipose package in Python.^40^ Erroneous segmentations were corrected with the Omnipose GUI or excluded from analysis. The resulting cell masks were used to calculate cell area and length (from the maximum Feret diameter), and for single-molecule analysis. Single-molecule detection, localization, and tracking were done with the SMALL-LABS algorithm.^41^ Single molecules were detected as non-overlapping spots with maximum pixel intensity values above the 95^th^ percentile pixel intensity of the frame. A single-molecule localization was performed by fitting the emission to a 2D Gaussian, and “good” single-molecule fits were defined as having a width no larger than 8 pixels (392 nm), a value of 2 for the SMALL-LABS stdtol parameter, and a value of 0.06 for the SMALL-LABS maxerr parameter for the maximum error on the fit. The positions were then connected into tracks by the Hungarian algorithm^42^ with a minimum track length of 4 steps and a maximum frame-to-frame step size of 15 pixels (735 nm).

Recorded trajectories were analyzed via a 2D mean squared displacement analysis (MSD) to calculate the apparent diffusion coefficient of that track, *D_app_*:

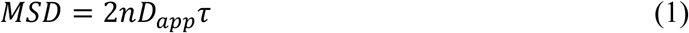

Where *n* = 2 is the dimensionality of the images, and *τ* is the time lapse. The camera integration time was 40 ms, and *D_app_* was calculated from *τ* = 40 – 160 ms using a weighted least-squares fit to Equation (1). Fits above a threshold of *R*^2^ ≥ 0.70 were used for further analysis (≥ 50% of the data), while others were excluded. The distribution of single-trajectory *D_app_* values was fit to a Gaussian mixture model to determine the average *D_app_* and weight fraction of each population.

#### Step-Size Analysis with Single-Molecule Analysis by Unsupervised Gibbs Sampling (SMAUG)

The same single-molecule trajectories were analyzed by the SMAUG algorithm, which uses a nonparametric Bayesian statistical framework to classify diffusive mobility states from trajectory data.^37^ SMAUG was run for 10,000 iterations on tracks with a minimum length of 6 steps and the initial number of states set to 5 to determine the number of mobility states, the average diffusion coefficient of each state, and the weight fraction of each state. A minimum of 45,000 steps was considered and analyzed for each condition.

#### Super-Resolution PALM Image Reconstruction

Super-resolution images of the nucleoid in each cell were generated by a Photoactivated Localization Microscopy (PALM) reconstruction.^43^ All drift-corrected HUα-PAmCherry localizations with a 95% confidence interval on the fitted position < 80 nm were selected. A Gaussian blur with a width equivalent to the 95% confidence interval on the fitted position was applied to each localization.

#### Nucleoid Occupancy Calculations from PALM Super-Resolution Images

The nucleoid occupancy was determined by fitting the distribution of pixel intensities in the PALM images with a two-component Gaussian mixture model. Pixel intensities below the fifth percentile and above the 99.95th percentile were excluded from analysis to remove outliers. A threshold pixel intensity value was set to 50% of the mean intensity of the second Gaussian distribution. Pixels with intensities greater than the set threshold were considered part of the nucleoid region.^24^ Nucleoid occupancy was calculated as the number of pixels in the nucleoid region divided by the number of pixels within the cell mask from cell segmentation of phase-contrast images via Omnipose as described previously.^24^

#### Calculation of Localization Heatmaps for Slow and Fast Trajectories

The coordinates of detected HUα-PAmCherry molecules and cell masks from Omnipose were used to generate the localization density heatmaps via the Python package *spideymaps*.^44^ For each cell, morphological skeletonization was used to define the midline bisecting the cell along the long axis. We plotted the localizations from slower molecules (trajectories with *D_app_ <* 0.15 µm^2^/s) separately from the localizations from faster molecules (trajectories with *D_app_* > 0.30 µm^2^/s). The cartesian coordinates (*x*, *y*) for each molecule were re-parameterized into a three coordinate system (*r*, *l*, *Φ*), where *r* is the radial distance from the defined midline, *l* is the longitudinal distance along the long axis of the cell found from projection onto the midline, and *Φ* is the angle measured relative to the midline in the pole regions of the cell. Parameters *r* and *l* were divided by the average cell width and total length, respectively, to determine the relative coordinates *r_rel_* and *l_rel_* such that all localizations share a common frame of reference.

Localizations were then binned based on *r_rel_*, *l_rel_*, and *Φ*. Because this approach results in unequal bin areas, bin areas were calculated for each bin in every cell. Localization counts and associated bin areas were summed across all cells. Areas and counts were symmetrized fourfold by adding counts and areas from symmetrically equivalent bins. The summed localizations were then divided by the summed areas to determine average localization density (counts per pixel).

### Bulk Fluorescence Microscopy

For exponential phase imaging, 1 mL of culture was removed and incubated with SYTOX green (500 nM) for 20 min. After incubation, cells were centrifuged at 5750 ×*g* for 2.5 minutes and then washed twice with spent medium warmed to 30 °C to remove unbound SYTOX green.

Cells were resuspended in 500 µL of spent medium for imaging. For stationary-phase cultures, 100 µL of cell culture was added to 900 µL of spent medium and stained with SYTOX green (10 µM). The mixture was then incubated for 45 minutes. The different protocols for exponential and stationary phase cell sample preparation were selected based on labeling efficiency to enable similar signal-to-noise ratios.

Phase-contrast images were collected on an Olympus IX-71 microscope with a 100× 1.45 NA phase-contrast oil-immersion objective with a 512 × 512-pixel EMCCD camera with a 100-ms exposure time. A CoolLED white light source with a 488-nm Chroma filter cube set was used to image stained DNA. The sample was illuminated alternatively with white light and 488-nm light to image the phase-contrast and fluorescence images of the nucleoid, respectively.

The cell boundary was determined from segmentation of the phase-contrast image by Omnipose, and the chromosome was visualized from SYTOX green fluorescence images. The nucleoid occupancy from nucleic acid staining was then determined from the number of pixels in the nucleoid area (as described previously^24^) divided by the number of pixels in the cell area.

### Real-Time Quantitative Reverse Transcription PCR (RT-qPCR)

Cell cultures were grown at 30 °C with shaking at 250 rpm and extracted at OD_600_ = 0.30 – 0.40 for exponential phase cells. Total RNA was extracted from cells using a Monarch Total RNA Miniprep Kit (NEB #T2010S) following miniprep instructions. Extracted RNA was quantified with a NanoDrop UV-Vis spectrophotometer by averaging three independent measurements. For RT-qPCR, a Luna Universal One-Step RT-qPCR kit (NEB #E3005L) was used. A master mix was prepared containing 1× Luna Universal One-Step Reaction Mix, 1× Luna WarmStart RT Enzyme Mix, 0.4 µM each of forward and reverse primer (for *dps*, *cysG*, or *hcaT*; **Table S1**), and nuclease-free water. For the RNA template, 200 ng of RNA was aliquoted into a 384-well plate and mixed with the master mix, then briefly centrifuged. For each RT-qPCR reaction, a no reverse transcriptase and a no-RNA template control were performed to control for DNA contamination. The RT-qPCR reactions were performed using a Bio-Rad CFX Opus 384 Real-Time PCR system. For RT-qPCR reactions, the following thermal profile was used: 55 °C for 10 min, 95 °C for 1 min, followed by 45 cycles of 95 °C for 10 s and 60 °C for 30 s. RT-qPCR curves were analyzed with the BR.io Bio-Rad software. The results were analyzed using the ΔΔ*C*_t_ method.^45^

## Results

### Dps-Mediated Nucleoid Compaction Slows HUα-PAmCherry Diffusion in Stationary Phase

We assessed how the dynamics of the NAP HUα depend on Dps by imaging HUα-PAmCherry expressed from the native HUα promoter in *E. coli* cells.^36^ HUα binds DNA without sequence specificity,^46^ and PAmCherry is a photo-activatable fluorescent protein that does not interfere with HUα function.^38^ We analyzed the positions of HUα-PAmCherry in WT and Δ*dps* cells growing at exponential and stationary phases by live-cell super-resolution (PALM) imaging (**Figure 1A)**, and we pooled the super-resolution localizations from all cells for each condition and plotted them as two-dimensional heatmaps (**Figure 1B**). Consistent with our previous findings, we found that the cell size depends on growth phase (**Figure 1B,C**) and the nucleoid is smaller in stationary-phase cells (**Figure 1B**).^24^ In exponential phase cells, the HUα-PAmCherry molecules are most abundant at the quarter cell positions. In contrast, in stationary phase, when the nucleoid is more compact, the HUα-PAmCherry localization pattern depends on Dps: in WT cells, HUα-PAmCherry molecules are concentrated evenly across the long axis of the nucleoid, while the HUα density is relatively enriched at the quarter positions in Δ*dps* cells (**Figure 1B**).

**Figure 1.**
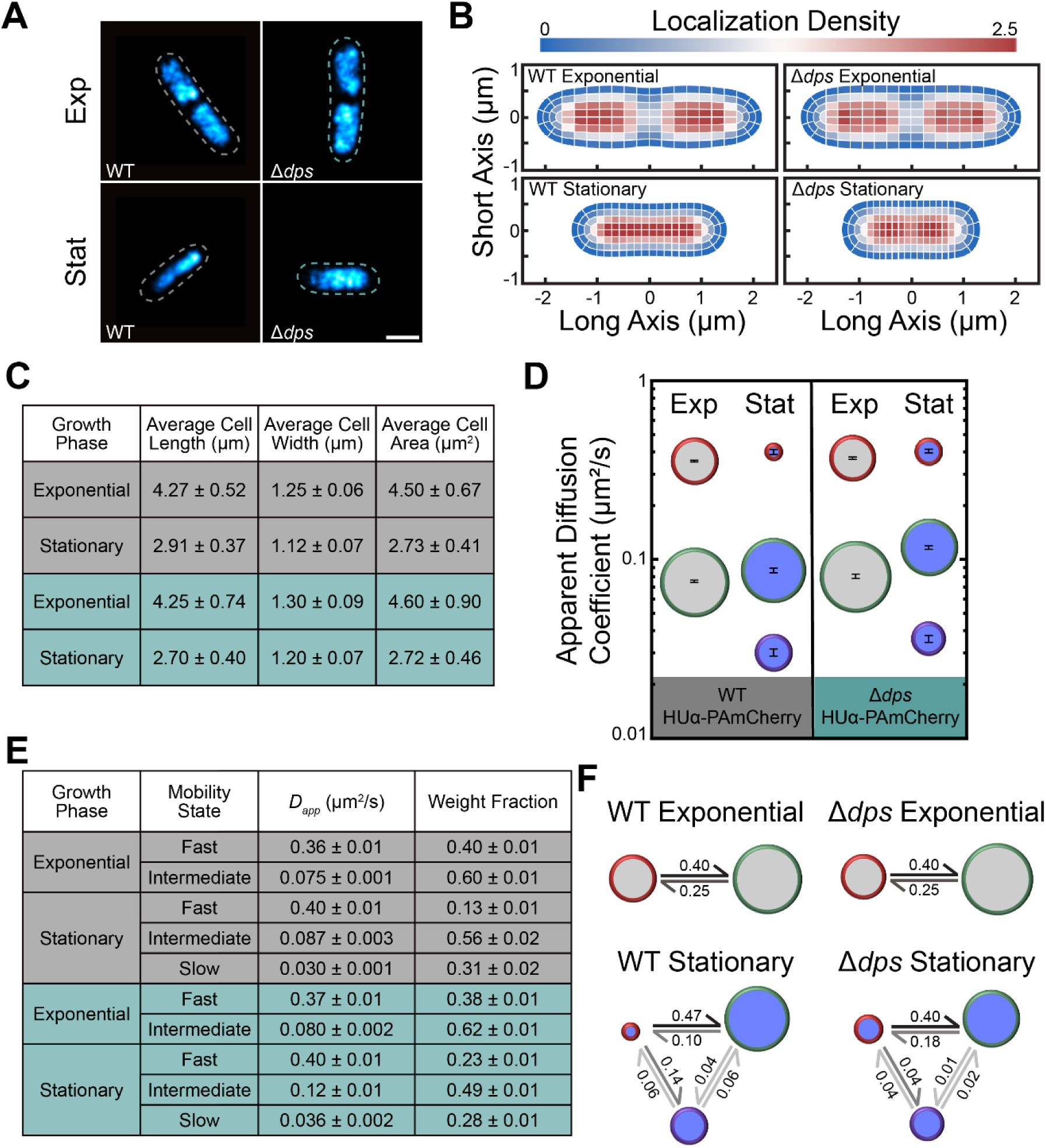
Analysis of HUα-PAmCherry Diffusion in WT and Δ*dps* Cells. **(A)** Super-resolution images of HUα-PAmCherry in representative WT and Δ*dps* cells in exponential and stationary phase. The dashed outlines depict the cell outlines from phase-contrast images of the cells. Scale bar: 1 µm. **(B)** Localization density heatmaps of HUα-PAmCherry localizations in WT and Δ*dps* cells in exponential and stationary phase. The lengths of the heatmaps along the short and long axes indicate the average cell length and width in the dataset, respectively. The colorscale denotes the localization density relative to the cell average. **(C)** Average cell lengths, widths, and areas of WT (grey) and Δ*dps* (green) HUα-PAmCherry cells used in the analysis. **(D) – (F)** Results from single-step diffusion analysis with SMAUG: **(D)** Average apparent diffusion coefficient for each diffusive state in WT and Δ*dps* cells in exponential (light grey) and stationary phase (dark blue). The diameter of each circle indicates that state’s weight fraction. The error bars denote the standard deviation of the iterations determining each average *D_app_*. **(E)** Calculated parameters for WT (grey) and Δ*dps* (green) HUα-PAmCherry cells. Error bars: standard deviation from 500 independent analysis iterations. **(F)** Probability of a molecule transitioning between each diffusive state in WT and Δ*dps* cells at exponential and stationary phases. The diameter of each circle indicates that state’s weight fraction. The probability of the transition is indicated next to the arrow.

We tracked single HUα-PAmCherry molecules in *E. coli* cells **(Figure S1A)**, and we assigned an average apparent diffusion coefficient, *D_app_*, to each HUα-PAmCherry trajectory based on the mean-squared displacement (MSD) (**Figure S1B**). Previous studies showed that Dps-mediated nucleoid compaction only occurred after 96 h of incubation.^24^ It is unclear why this compaction does not happen at an earlier time point (e.g., 24 h). Nonetheless, we use the 96-h timepoint as deep stationary phase in the current study. In exponential phase, we observed no difference in the *D_app_* distributions for WT and Δ*dps* cells (**Figure S1C**), which is consistent with low Dps expression in exponential phase.^19^ However, the *D_app_* distribution for WT cells is slower in stationary phase relative to exponential phase (**Figure S1D**), which is correlated with increased nucleoid compaction in stationary phase.^24^ Additionally, the *D_app_* distribution is similar for Δ*dps* cells in exponential and stationary phase, indicating that Dps-mediated nucleoid compaction is at least partly responsible for the slowed HUα diffusion seen in stationary phase in WT cells. It is also possible that Dps and HUα directly interact, which would slow HUα diffusion.^6^

To determine the spatial patterning of the slowest-moving HUα-PAmCherry molecules (corresponding to HUα that is either bound to or frequently interacting with DNA), we pooled the super-resolution localizations from only the slower trajectories (*D_app_* < 0.15 µm^2^/s) for all cells for each condition and plotted them as two-dimensional heatmaps (**Figures S2, S3**). In exponential phase, slow-moving HUα molecules are most often found at the cell quarter positions, consistent with the more densely packed DNA slowing HUα with frequent interactions. However, fast-moving molecules (*D_app_* > 0.30 µm^2^/s) are more enriched at the mid-cell, consistent with a higher DNA density at the quarter positions. Overall, there is no notable difference in the localization patterns of the fast or slow molecules between exponential-phase WT and Δ*dps* cells (**Figure S2**). On the other hand, in stationary phase, the HUα localization pattern is Dps-dependent (**Figure 1B**), and we find that the enrichment of HUα at the quarter positions in stationary-phase Δ*dps* cells corresponds specifically to an enrichment of slow-moving HUα-PAmCherry molecules at these positions (**Figure S3**).

### Analyzing the Distribution of Steps Within the Heterogeneous Single-Molecule Trajectories Indicates Three Distinct Types of HUα-PAmCherry Motion in the Stationary Phase

The single-trajectory MSD analysis provides only the average *D_app_* of each trajectory, but target binding and changes in biological function can cause a change in a molecule’s diffusive behavior in the middle of a trajectory.^37^ In addition, we measured asymmetric *D_app_* distributions for all conditions (**Figure S1C,D**); this asymmetry indicates multiple underlying distributions. To capture the heterogeneities within single trajectories, we used the nonparametric Bayesian statistics algorithm SMAUG to analyze the HUα-PAmCherry trajectory dataset. For each dataset, we inferred the mobility states within the distribution of *s* = 50,000 – 100,000 step sizes rather than grouping them into trajectories.^37^

Analysis of HUα-PAmCherry diffusion in WT exponential phase cells inferred two mobility states, one fast and one intermediate (**Figure 1D,E**). The high transition probability between these two states indicates that HUα switches from the intermediate state to the fast state and back on the scale of the 40-ms image frame time (**Figure 1F**). These results for HUα-PAmCherry diffusion in exponential phase are similar to previous findings by Bettridge et al.,^38^ who also found that HUα rapidly switches between a slower and a faster mobility state; because they found that HUα-PAmCherry had similar confinement and diffusion coefficient to that of a stably bound TetR-mCherry protein, Bettridge et al. inferred that the slower moving molecules were in a bound state with movement that was the result of only intrinsic chromosomal DNA fluctuations. In WT cells, the weight fraction of the fastest diffusive state decreases from 40% in exponential phase to 13% in stationary phase, and a third, slower mobility state that comprises 31% of the HUα-PAmCherry molecules is inferred in stationary phase with 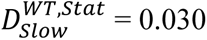 µm^2^/s (**Figure 1D,E**). This slowest mobility state is, in fact, even slower than the “bound” state observed in exponential phase by Bettridge et al.,^38^ indicating that either the nucleoid is much more densely packed in stationary phase or the slow HUα molecules in exponential phase interact very frequently with DNA rather than being bound. Furthermore, we deduced that HUα is stably bound or extremely confined in this slowest state based on the high probability of transitioning <u>into</u> the slowest state from the fastest (0.14 probability) relative to the very low probabilities of transitioning <u>out</u> of the slow state to either of the other states.

### Dps Slows HUα-PAmCherry Diffusion in Stationary Phase

The number of HUα-PAmCherry mobility states, the average *D_app_* of those states, and the transition probabilities do not depend on Dps in exponential phase (**Figure 1F**), as expected given the very low Dps expression levels in the exponential phase.^19^ However, in stationary phase, the distribution of single-molecule HUα-PAmCherry diffusion coefficients significantly depend on Dps (**Figure S1D**). This dependence may be attributed to the subtly decreased nucleoid compaction in Δ*dps* cells^24^ or to other factors, including possible interactions between HUα and Dps.^6^ Furthermore, the single-step analysis indicates that the increased mobility of HUα in stationary-phase Δ*dps* cells is due to a large increase in the weight fraction of the fastest diffusive state (from 13% in WT cells to 23% in Δ*dps* cells), and a decrease in the weight fraction and an increase in the average *D_app_* for the slowest HUα mobility state in Δ*dps* cells relative to WT cells. Most prominently, we measured an increased average *D_app_* for the intermediate state (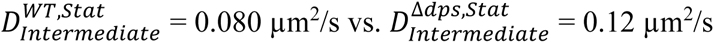; **Figure 1D,E**).

Furthermore, the probability of transitioning from the intermediate state to the fast state nearly doubles from 0.10 in WT cells to 0.18 in Δ*dps* cells (**Figure 1F**). Concomitantly, there is a greatly reduced probability of transitioning from the intermediate to slow state (probability = 0.01), and the transition probability from the fast state to slow state is reduced in Δ*dps* cells compared to WT.

This increased HUα diffusivity, increased transition probabilities to the fastest state, and reduced transition probabilities to the slow state in stationary phase Δ*dps* cells may be attributed to fewer regions of very dense DNA in the less compact nucleoid, which correlates well with the lower localization densities of HUα in stationary phase Δ*dps* cells compared to WT cells **(Figure 1B)**. Overall, Dps, which subtly compacts the dense nucleoid in stationary phase, has the effect of reducing the dynamics of HUα.

### Nucleoid Compaction Increases in Exponential-Phase Δ*hns* Cells

To complement our investigation of the prevalent stationary-phase NAP, Dps, we investigated how the abundant exponential-phase NAP, H-NS, affects nucleoid structure and HUα-PAmCherry diffusion. Representative fluorescence images of the nucleoid labeled with the nucleic acid stain SYTOX Green in WT and Δ*hns* cells in exponential and stationary phase are shown in **Figure 2A**. 48 h after inoculation of cells into fresh HDA medium at 30 °C, the culture density saturates at a lower OD_600_ value in the Δ*hns* cell culture than in WT (**Figure 2B**). This result indicates that H-NS provides a fitness advantage for cells in stationary phase. Although the mechanism of how H-NS protects the cell is unclear, our results are consistent with previous reports by Chib and Mahadevan for *E. coli* strains with *hns* mutations.^47^ The mean lengths of WT and Δ*hns* HUα-PAmCherry cells at exponential phase (mean ± standard deviation) are 3.76 ± 0.81 µm and 3.99 ± 0.85 µm, respectively. In stationary phase, Δ*hns* HUα-PAmCherry cells are much longer on average: the mean lengths of WT and Δ*hns* HUα-PAmCherry cells are 2.07 ± 0.45 µm and 2.71 ± 0.57 µm, respectively. Thus, we find that the deletion of H-NS significantly impacts the cell length in stationary phase, and to a lesser extent in exponential phase (**Figure 2C,D**).

**Figure 2.**
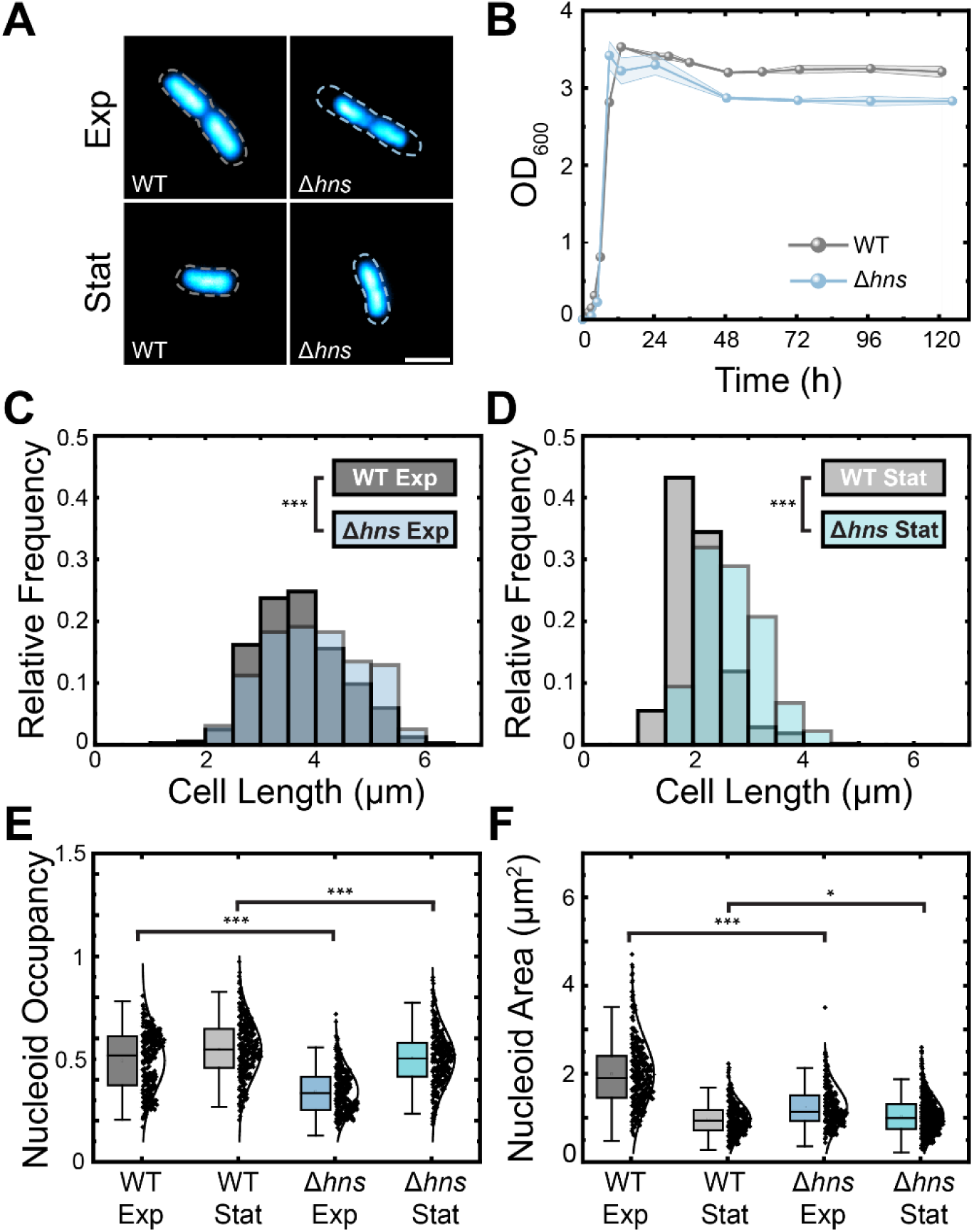
Characterization of Δ*hns* HUα-PAmCherry cells. **(A)** Bulk fluorescence images of SYTOX Green-stained nucleoids in representative WT and Δ*hns* cells in exponential phase and stationary phase. The dashed outlines depict the cell outlines, taken from phase-contrast images of the cells. Scale bar: 2 µm. **(B)** Growth curves for WT and Δ*hns* HUα-PAmCherry cells. Shading indicates the standard deviation from 3 replicates. **(C), (D)** Distributions of WT and Δ*hns* cell lengths for exponential and stationary phase cells. Statistical significance determined through a two-tailed t-test (***: *p* < 0.0001). **(E)** Nucleoid occupancy box and whisker plots for WT and Δ*hns* cells in exponential and stationary phase. Same color designations as in (C) and (D). Whiskers denote two standard deviations about the mean. The center line is the median of the dataset. Statistical significance determined through a two-tailed t-test (***: *p* < 0.0001), (*: *p* < 0.01). **(F)** Same as (E), but for nucleoid area distributions.

Next, we investigated the effects of H-NS on nucleoid compaction based on the nucleoid occupancy (the ratio of the area of the SYTOX Green stain to the area of the cell) in exponential and stationary phase cells (*n* = 300 each, **Figure 2E**). The nucleoid occupancy is most different between WT and Δ*hns* cells in exponential phase, with average nucleoid occupancies of (mean ± 95% confidence interval) 0.49 ± 0.02 and 0.34 ± 0.01, respectively, indicating that the nucleoid is more compact in exponential-phase Δ*hns* HUα-PAmCherry cells. We also measured a lower nucleoid occupancy in exponential phase for Δ*hns* compared to WT HUα-PAmCherry cells from super-resolution maps of HUα-PAmCherry (**Figure S4**). On the other hand, in stationary phase, Δ*hns* cells have only slightly smaller nucleoid occupancy (0.55 ± 0.02 and 0.50 ± 0.02, respectively, for WT and Δ*hns* cells). Because the change in the nucleoid occupancy is partly due to a change in the cell length (**Figure 2C,D**), we compared the nucleoid areas as well (**Figure 2F**). In exponential phase, the Δ*hns* cells have a much smaller nucleoid relative to the WT (average nucleoid areas of 2.00 ± 0.09 µm^2^ and 1.24 ± 0.05 µm^2^, respectively, for WT and Δ*hns* cells). However, in stationary phase, the average nucleoid areas are more comparable for WT and Δ*hns* HUα-PAmCherry cells (0.98 ± 0.04 µm^2^ and 1.15 ± 0.05 µm^2^, respectively) which indicates that the measured changes in nucleoid occupancy are in fact due to changes in the cell areas.

### H-NS Represses *dps* Transcript Levels

Given that H-NS has been proposed to play a role in nucleoid compaction^27,35,48,49^ by promoting DNA bridging, it is somewhat surprising that the nucleoids of Δ*hns* cells are more compacted than those of WT cells in the exponential phase (**Figure 2E,F**). We therefore considered that H-NS is also a well-known global gene silencer^28,35,50,51^ that is known to selectively repress *dps* expression in the exponential phase.^52^ We evaluated the effect of H-NS on Dps expression in exponential phase by real-time quantitative PCR (RT-qPCR), and we confirmed that transcript levels increase in Δ*hns* cells (**Figure S5**).^53^ However, because the little Dps present in exponential phase cells is rapidly degraded,^19,54^ we expect H-NS to have only a mild impact on Dps concentration in exponential phase. Instead, it is likely that one of the other NAPs regulated by H-NS alters nucleoid compaction in the exponential phase.

### Deleting H-NS Leads to a Less Compact Exponential-Phase Nucleoid

We super-resolved the HUα-PAmCherry positions in a set of *n* = 66 WT cells and a set of *n* = 40 Δ*hns* cells in exponential phase, selecting cells with similar lengths (∼4 µm) in both strains for a balanced comparison, and we pooled the super-resolution localizations from all cells for each condition and plotted them as two-dimensional heatmaps (**Figure 3A-C**). We also compared *n* = 44 WT cells and *n* = 36 Δ*hns* cells in stationary phase, with lengths ∼2.8 µm (**Figure 3A-C**). We found that deleting H-NS compacts the exponential-phase HUα localization pattern and has little effect on the size of the stationary-phase nucleoid.

**Figure 3.**
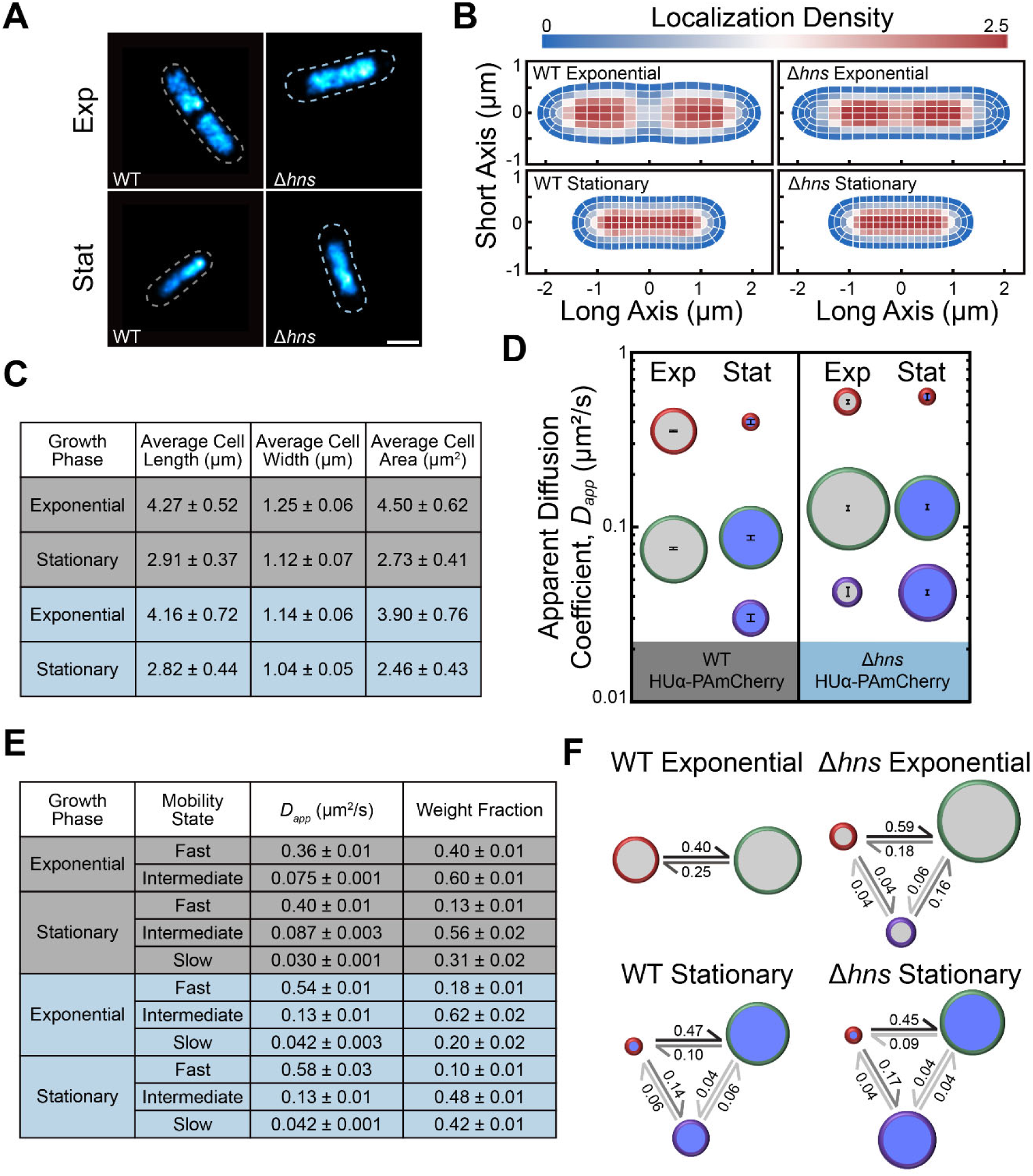
Analysis of HUα-PAmCherry Diffusion in Δ*hns* cells. **(A)** Representative super-resolution images of HUα-PAmCherry in WT and Δ*hns* cells in exponential and stationary phase. The dashed outlines depict the outline of the cell, taken from phase-contrast images of the cell. Scale bar: 1 µm. **(B)** Density heatmaps of HUα-PAmCherry localizations in Δ*hns* cells in exponential and stationary phase. The lengths of the heatmaps along the short and long axes indicate the average cell length and width in the dataset, respectively. The colorscale denotes the localization density relative to the cell average. **(C)** Average cell lengths, widths, and areas of WT (grey) and Δ*hns* (blue) HUα-PAmCherry cells used in the analysis. **(D) – (F)** Results from single-step diffusion analysis with SMAUG: **(D)** Average apparent diffusion coefficient for each diffusive state in WT and Δ*dps* cells in exponential (light grey) and stationary phase (dark blue). The diameter of each circle indicates that state’s weight fraction. The error bars denote the standard deviation of the iterations determining each average *D_app_*. **(E)** Calculated parameters for WT (grey) and Δ*hns* (blue) HUα-PAmCherry cells. Error bars: standard deviation from 500 independent analysis iterations. **(F)** Probability of a molecule transitioning between each diffusive state in WT and Δ*hns* cells at exponential and stationary phases. The diameter of each circle indicates that state’s weight fraction. The probability of the transition is indicated next to the arrow.

### H-NS Slows HUα Diffusion in Exponential Phase

To determine if nucleoid decompaction by H-NS in exponential phase correlates with increased HUα-PAmCherry diffusion, we tracked HUα-PAmCherry in Δ*hns* cells. Surprisingly, though we measured increased nucleoid compaction in the Δ*hns* cells relative to the WT cells (**Figure 2E)**, we measured similar HUα diffusion upon H-NS deletion in exponential-phase cells (**Figure S6A**). We measured slower diffusion in the average HUα diffusion upon H-NS deletion in stationary-phase cells (**Figure S6B**).

We analyzed the single-step displacements with SMAUG Bayesian analysis. In exponential phase, the average *D_app_* for the fast and intermediate HUα-PAmCherry mobilities states is greater in Δ*hns* cells than in the WT background, though deleting H-NS leads to the appearance of a third, very slow mobility state (**Figure 3D,E**). While these slowest-moving HUα-PAmCherry molecules indicate a population of molecules bound to DNA, the probability of transitioning from this slowest state to the intermediate, transiently interacting state, is high (0.16) in Δ*hns* cells (**Figure 3F**), which indicates that, while dense regions of DNA exist, HUα-PAmCherry easily moves throughout the nucleoid without spending much time confined by the nucleoid. Concomitantly, the slow HUα trajectories are less confined to the quarter positions in Δ*hns* cells (**Figure S7**) than in WT cells (**Figure S2**).

On the other hand, the average *D_app_* values for the three HUα-PAmCherry mobility states are the same in exponential and stationary phase Δ*hns* cells (**Figure 3D,E**). However, the associated weight fractions change: at stationary phase, the slowest mobility state is very prominent (42%) in Δ*hns* cells; this state has only a 20% weight fraction in Δ*hns* exponential-phase cells and has a 31% weight fraction in the WT stationary-phase cells. The stationary-phase nucleoid organization is therefore different in Δ*hns* cells relative to WT cells, and the increased fraction of slow-moving HUα molecules suggests increased regions of dense DNA in Δ*hns* cells. These observations indicate that H-NS plays a large role in the maintenance of the nucleoid architecture across growth phases, even though the H-NS concentration decreases in stationary phase.^19^ **<u>Discussion</u>**

We investigated how NAPs affect one another’s binding and diffusion, and hence nucleoid organization, based on measuring the diffusion of the non-specific DNA-binding NAP HUα-PAmCherry. Consistent with previous work,^38^ we found that HUα-PAmCherry can easily transition between two distinct mobility states in exponential phase (**Figure 1C**). These two states were previously considered to be a stably DNA-bound HUα state and a state in which HUα transiently binds to DNA for a short time before releasing and then binding again to nearby DNA.^38^ Overall, our results are consistent with a model in which DNA is relatively accessible in exponential phase and, therefore, the slow state in exponential phase is consistent with HUα frequently interacting and binding to less densely packed and more mobile DNA.

In stationary-phase cells, we detected a third, slower mobility state for HUα, consistent with reports of regions of increased DNA compaction in stationary phase and increased DNA confinement.^8,10,24,55–58^ This slowest diffusive state may be attributed to the confinement of HUα to very crowded regions of the nucleoid and/or more stable binding to DNA. We found that a significant percentage of HUα-PAmCherry molecules move very slowly in stationary phase (**Figure 1D**), which is consistent with nucleoid compaction:^19,58^. Our data supports the interpretation that the slowest mobility state describes molecules confined within the compact nucleoid, as the transition probabilities for leaving the third mobility state are extremely low (< 0.06; **Figure 1F**), indicating trapping in dense regions of DNA.

How HUα-PAmCherry interacts with DNA depends on the environment. In exponential phase, DNA is less densely packed and more mobile, which we hypothesize leads to two populations: a slow interacting population that is either trapped or stably bound by DNA, and a fast, frequently interacting population. Previous work by Bettridge et al. measured a stably bound slow population, and also reported 100-ms dwell times,^38^ in line with HUα-PAmCherry being trapped by DNA. On the other hand, the nucleoid is more compacted in the stationary phase, and we find an additional, slower population of HUα-PAmCherry molecules. This new mobility state could result either from HUα-PAmCherry being bound to DNA with much longer dwell times, or from the chromosomal DNA being much less dynamic compared to the exponential phase.^38^

We measured the effects of Dps on HUα-PAmCherry mobility and observed no impact of Dps on HUα-PAmCherry diffusion in exponential phase. However, we observed subtle Dps-mediated nucleoid compaction in the deep stationary phase (96 h), which coincides with slower HUα-PAmCherry motion.

We investigated the effects of nucleoid organization by H-NS. We found that H-NS provides a fitness advantage in stationary phase (**Figure 2B**)^47,59,60^ and that the average stationary-phase cell length is much longer in Δ*hns* cells than in WT cells. Surprisingly, we found that the nucleoid is more compacted in exponential phase Δ*hns* cells than in WT cells, while the nucleoid is slightly larger in Δ*hns* stationary phase cells relative to WT (**Figure 2E,F**). Therefore, H-NS differently impacts nucleoid size in exponential and stationary phase.^19^ Although H-NS may have pleiotropic effects on gene regulation and global nucleoid organization, it is clear that H-NS greatly contributes to proper nucleoid organization in exponential phase.

We investigated how changes in nucleoid organization by H-NS correlate with HUα-PAmCherry dynamics. We found that H-NS decreases HUα-PAmCherry mobility in exponential phase (**Figure 3D,E**). In addition, the probability is very low for molecules in the fast or intermediate states to transition to the slowest state in Δ*hns* cells. Furthermore, we observed a third, very slow state in exponential-phase Δ*hns* cells, consistent with strongly confined HUα-PAmCherry. This slow state suggests that deleting H-NS increases DNA density in certain regions within the nucleoid. Overall, these results show that H-NS-mediated nucleoid organization is important in exponential phase. Finally, we examined how H-NS impacts HUα-PAmCherry diffusion in stationary phase, which revealed three distinct HUα-PAmCherry mobility states in Δ*hns* cells with the same average *D_app_* values as in exponential phase Δ*hns* cells. The transition probabilities also changed in stationary phase. Compared to WT cells, the Δ*hns* cells have slightly less compacted nucleoids in stationary phase, which may have caused faster HUα-PAmCherry diffusion. Alternatively, H-NS is known to regulate DNA topology for gene silencing as well as for structural maintenance of the nucleoid,^27,49,61^ which may affect HUα access and mobility.

## Conclusions

Overall, our findings show that Dps-mediated nucleoid compaction correlates with overall slower HUα-PAmCherry diffusion and a third population of very slow-moving HUα molecules in stationary phase. Additionally, we determined that H-NS affects the nucleoid size and impacts HUα diffusion in both exponential and stationary phases, resulting in a significant population of stably bound HUα state in both growth phases. Overall, we determined that NAPs not only regulate the nucleoid microenvironment, but they also influence how other NAPs interact with DNA, consistent with the model that NAPs cooperate to maintain and structure the nucleoid.

## Supporting Information

Supplemental Figures S1 – S7, Supplemental Table S1.

## Acknowledgements

The HUα-PAmCherry strain was a generous gift from Xiaowei Zhuang. Thanks to Dr. Daniel Foust for providing the Spideymaps heatmaps code. Support for this work came primarily from the National Institutes of Health Grant R01 GM143182 to J.S.B. and X.W. L.A.M. gratefully acknowledges NIH training grant K12GM111725. Additional support came from National Institutes of Health R01GM141242 (X.W.), R01AI172822 (X.W.), and R01GM144731 (J.S.B.). This research is a contribution of the GEMS Biology Integration Institute, funded by the National Science Foundation DBI Biology Integration Institutes Program, Award #2022049 (X.W.).

