## Supplemental Information for "Characterizing the Impact of Nucleoid-Associated Proteins on HU-DNA Interactions by Live-Cell Single-Molecule Tracking"

**SUPPORTING INFORMATION FOR:**

### Characterizing Dynamic Chromosome Organization by Nucleoid-Associated Proteins Dps and H-NS by Live-Cell Single-Molecule Tracking

Fuller, D. E. H.[1], Dai, X.[1], McCarthy, L. A.[1], Way, L. E.[2], Wang, X.[2], and Biteen, J. S.[1,*]

[1] Department of Chemistry, University of Michigan, Ann Arbor, MI, 48109, USA

[2] Department of Biology, Indiana University, Bloomington, IN, 47405, USA

**CONTENTS:**

- Supplemental Figures S1 – S7
- Supplemental Table S1

#### SUPPLEMENTAL FIGURES

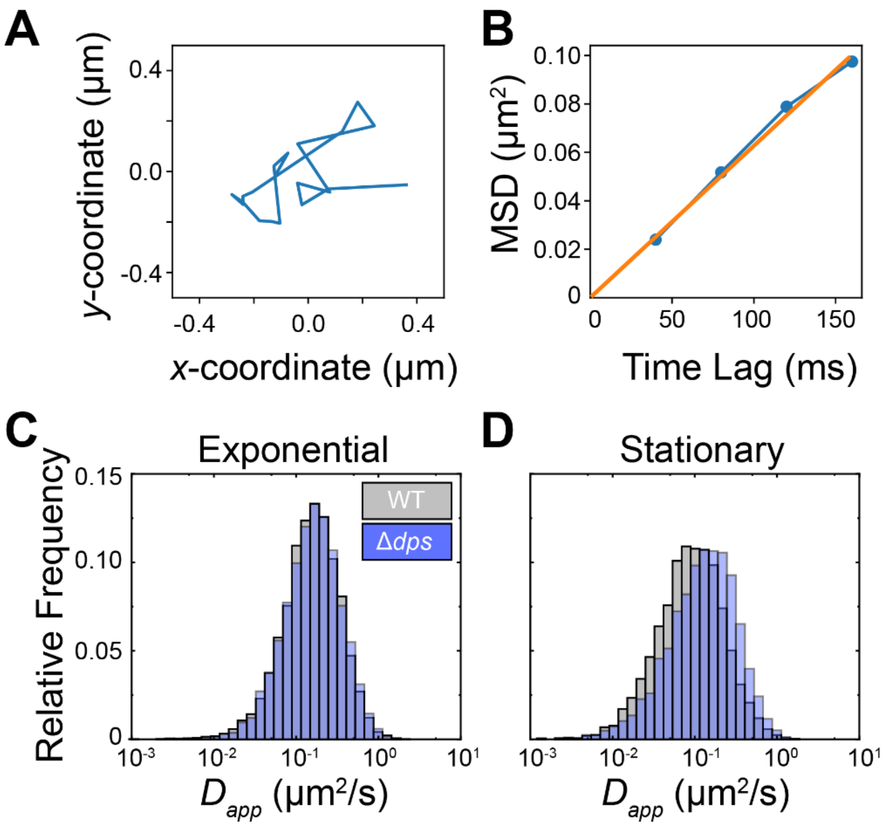


**Figure S1. MSD analysis of wild-type and Δ*dps* HUα-PAmCherry cells. (A)** Representative trajectory of HUα-PAmCherry. The single-molecule position is recorded every 40 ms. **(B)** Mean squared displacement (blue) and fit (orange) for the data in the trajectory shown in **(A)**. **(C, D)** Distributions of the apparent diffusion coefficient, $D_{app}$, for $n$ = 4153 – 7554 single-molecule trajectories in $N$ = 44 – 62 cells for **(C)** WT (grey) and Δ*dps* (blue) cells in exponential phase, and **(D)** WT and Δ*dps* cells in stationary phase.

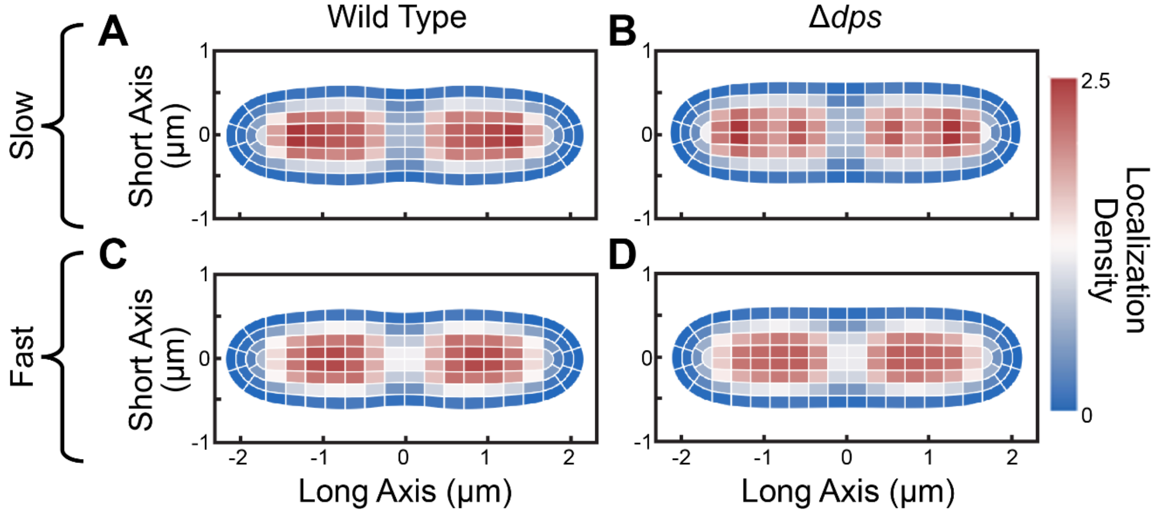


**Figure S2. Localization density heatmaps of HUα-PAmCherry molecules in exponential-phase cells. (A), (B)** Localizations of molecules in slow trajectories ($D_{app}$ < 0.15 μm$^2$/s) in **(A)** WT cells and **(B)** Δ*dps* cells. **(C), (D)** Localizations of molecules in fast trajectories ($D_{app}$ > 0.30 μm$^2$/s) in **(C)** WT cells and **(D)** Δ*dps* cells. The colorscale denotes the localization density relative to the cell average.

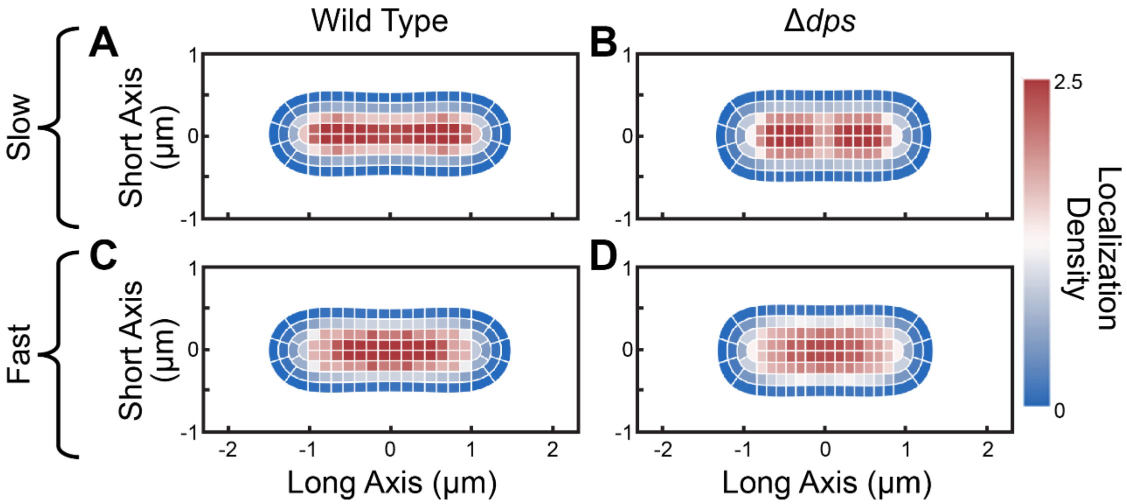


**Figure S3. Localization density heatmaps of HUα-PAmCherry molecules in stationary-phase cells.** **(A), (B)** Localizations of molecules in slow trajectories ($D_{app}$ < 0.15 μm$^2$/s) in **(A)** WT cells and **(B)** Δ*dps* cells. **(C), (D)** Localizations of molecules in fast trajectories ($D_{app}$ > 0.30 μm$^2$/s) in **(C)** WT cells and **(D)** Δ*dps* cells. The colorscale denotes the localization density relative to the cell average.

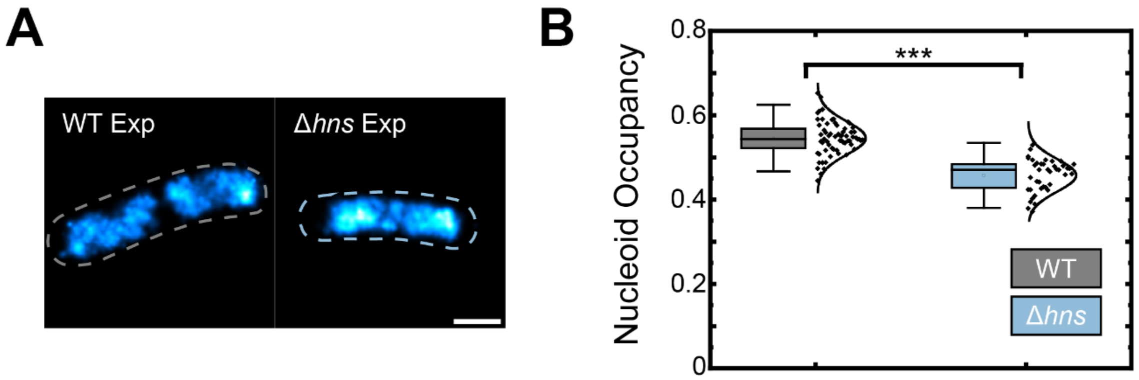


**Figure S4. Super-resolution analysis of WT and Δ*hns* HUα-PAmCherry cells in the exponential phase.** **(A)** Representative super-resolution images of exponential phase WT and Δ*hns* HUα-PAmCherry cells. Scale bar: 1 µm. **(B)** Nucleoid occupancy distributions determined from super-resolved nucleoids based on imaging HUα-PAmCherry in WT and Δ*hns* cells. Whiskers denote two standard deviations about the mean, and the center line of the box indicates the median of the dataset. Statistical significance determined through a two-tailed t-test (***: $p < 0.0001$).

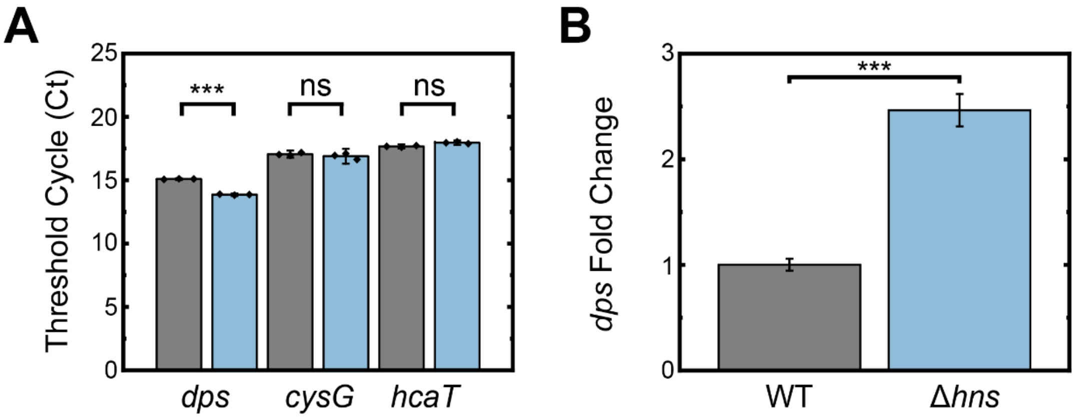


**Figure S5. Real-time quantitative PCR of *dps* transcripts in exponential phase WT and Δ*hns* HUα-PAmCherry cells. (A)** Plots of threshold cycle (*Ct*) values for *dps*, *cysG*, and *hcaT* between exponential phase WT and Δ*hns* HUα-PAmCherry cells. Error bars indicate the standard deviation of three biological replicates. Grey: WT HUα-PAmCherry cells, blue: Δ*hns* HUα-PAmCherry cells. **(B)** Fold expression change of *dps* transcripts in exponential phase WT and Δ*hns* HUα-PAmCherry cells, as determined by the $2^{-\Delta\Delta Ct}$ method. Error bars indicate the standard deviation of three biological replicates. Statistical significance determined through a two-tailed t-test (***: $p < 0.0001$; ns: $p > 0.05$).

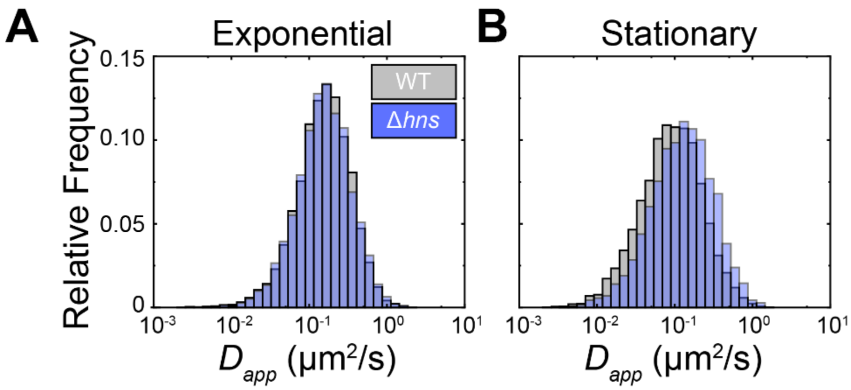


**Figure S6. MSD analysis of wild type vs Δ*hns* HUα-PAmCherry cells. (A, B)** Distributions of apparent diffusion coefficient, $D_{app}$, for $n$ = 2804 – 7554 single-molecule trajectories in $N$ = 37 – 62 cells for **(A)** WT (grey) and Δ*hns* (blue) cells in exponential phase, and **(B)** WT and Δ*hns* cells in stationary phase.

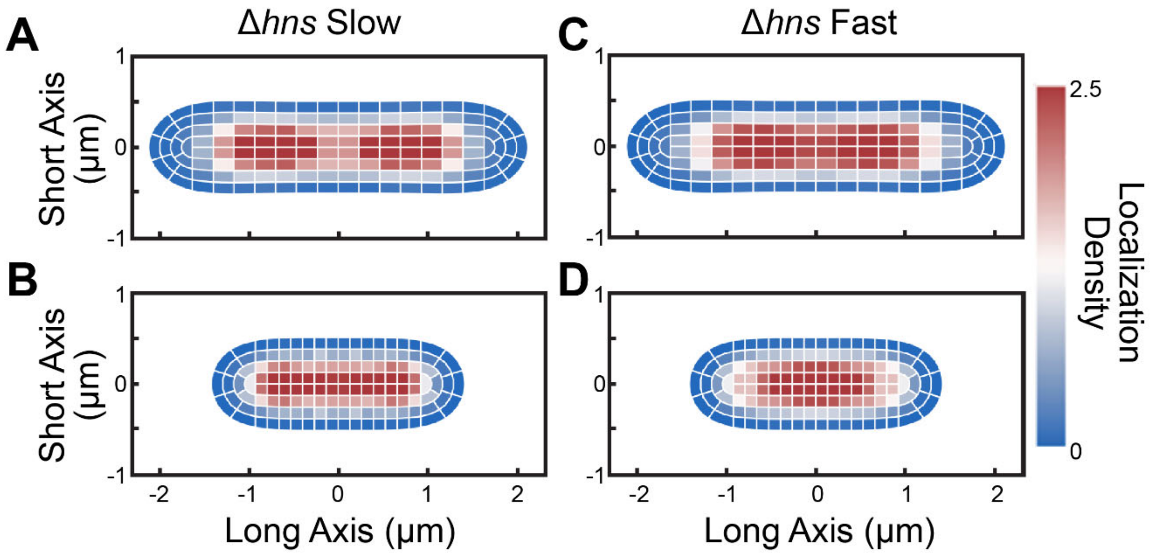


**Figure S7. Localization density heatmaps of HUα-PAmCherry molecules within slow ($D_{app}$ < 0.15 µm$^2$/s) and fast trajectories ($D_{app}$ > 0.30 µm$^2$/s. (A)** Localizations of slow molecules in exponential phase Δ*hns* cells. **(B)** Localizations of slow molecules in stationary phase Δ*hns* cells. **(C)** and **(D)** Same as (A) and (B), but for fast Δ*hns* cells. The lengths of the heatmaps along the short and long axes indicate the average cell length and width in the dataset, respectively. The colorscale denotes the localization density relative to the cell average.

**SUPPLEMENTAL TABLE**

**Table S1**. Forward and reverse RT-qPCR primers.

| **Oligonucleotide Name** | **Sequence (5′ to 3′)** |
|---|---|
| *dps* forward | GTCGTCATCTTTCGCTTCGC |
| *dps* reverse | ACCCCGCTGAAAAGTTACCC |
| *cysG* forward | CCAGCGTCTGTTTTTCTGCC |
| *cysG* reverse | TTCGGGTATTCCACTCACGC |
| *hcaT* forward | AACGGATGACTTCGCTACCC |
| *hcaT* reverse | GTTGCCGTGGTTGATAGTGG |
